# Comparative Metabolomic Profiling Reveals Salinity Tolerance Mechanisms in a Rice Introgression Line

**DOI:** 10.64898/2026.07.06.736799

**Authors:** Chander Kant Chaudhary, Praveen Kumar Guttula, Kirti Agrawal, Prasanta Kumar Subudhi, Manas Ranjan Gartia

**Affiliations:** School of Plant, Environmental, and Soil Sciences, Louisiana State University Agricultural Center, Baton Rouge, LA 70803, USA; Department of Mechanical and Industrial Engineering, Louisiana State University, Baton Rouge, LA 70803, USA

**Author notes:** Corresponding authors: Manas Ranjan Gartia, Department of Mechanical and Industrial Engineering, Louisiana State University, Baton Rouge, LA 70803, USA, Prasanta Kumar Subudhi, School of Plant, Environmental, and Soil Sciences, Louisiana State University Agricultural Center, Baton Rouge, LA 70803, USA.

**Keywords:** Rice, Salinity tolerance, Untargeted metabolomics, Differentially accumulated metabolites, Redox homeostasis, Amino acid metabolism, Abscisic acid

## Abstract

Rice (*Oryza sativa*) is highly sensitive to salinity, yet the metabolic mechanisms underlying salt tolerance remains incompletely understood. In this study, we performed leaf tissue-specific untargeted metabolomic profiling of the salt-tolerant introgression line JN100 (JN), its donor parent Nona Bokra (NB), and its recurrent parent Jupiter (JU) to characterize metabolic responses to salt stress. Comparative analysis identified differentially accumulated metabolites (DAMs) spanning diverse chemical classes, including amino acids, sugars and carbohydrates, lipids, organic acids, cofactors, electron carriers, and nucleotides. Under salt stress (SS), 201 DAMs (89 upregulated and 112 downregulated) were detected in JN relative to JU. Notably, metabolites such as allantoin, glycitin, nicotinamide ribotide, D-arabinono-1,4-lactone, violanthin, L-methionine S-oxide, ribitol, lysine, rutin, glutamine, pantothenic acid, and quinic acid, showed significant differential accumulation. Pathway enrichment analysis revealed significant enrichment of arginine biosynthesis, purine metabolism, and alanine, aspartate, and glutamate metabolism, indicating extensive reprogramming of nitrogen and energy-associated metabolic pathways under salinity stress. Integration of transcriptomic and metabolomic datasets from the SS experiments further identified ten differentially expressed genes (DEGs) associated with the metabolite network in the JN vs. JU comparison. Among these, *OsDHQDT/SDH*, *OsFd-GOGAT*, *phenylalanyl-tRNA synthetase*, *OsP5CS1*, *OsP5CS2*, and a *pyridoxal phosphate-dependent transferase* were linked to metabolites involved in shikimate, amino acid, and proline metabolism. Collectively, these results demonstrate that salinity tolerance in rice is associated with coordinated transcriptional and metabolic reprogramming that supports oxidative stress mitigation and adaptive stress responses.

## Introduction

Plants have evolved adaptive mechanisms to withstand diverse abiotic stresses, including soil salinity, drought, and high temperatures, which adversely affect growth, development, and productivity. Rice, a glycophyte, is highly sensitive to salt stress (Hoang et al. 2016), with a tolerance threshold of only about 3 dS/m, whereas soil is classified as saline at levels above 4 dS/m ((Tamanna et al. 2024; Singh et al. 2021). Metabolites such as proline and trehalose accumulate under saline conditions in plants and act as key modulators and phenotypic effectors (Ashraf and Foolad 2007). Salinity primarily causes osmotic and ionic imbalances in plant cells, ultimately leading to severe growth inhibition and even plant death. The initial osmotic stress triggered by salt exposure is gradually followed by an ionic stress phase as toxic ions accumulate within plant tissues. Rice landraces cultivated in coastal regions are frequently exposed to saline conditions. Among them, several traditional and wild landraces exhibit salinity tolerance; however, this tolerance is often associated with reduced grain yield (Ishikawa et al. 2022). For instance, the wild rice *Oryza coarctata* exhibits exceptional salinity tolerance and can withstand salt concentrations greater than 40 dS/m (Ishikawa et al. 2022). Under saline conditions, rice plants undergo severe ionic imbalance, characterized by excessive accumulation of Na and Cl ions, which disrupts vital physiological processes, including photosynthesis, protein synthesis, and enzyme activity (Radwan et al. 2026).

Abiotic stress conditions induce excessive production of reactive oxygen species (ROS), leading to oxidative damage to proteins, DNA, and membrane lipids. Metabolites such as ascorbic acid, glutathione, flavonoids, carotenoids, and proline help scavenge these ROS and protect cells from oxidative stress (Dong et al. 2020; Nakabayashi and Saito, 2015). Flavonoids and glutathione function as antioxidants under abiotic stress conditions and minimize the oxidative damage (Dong et al. 2020; Nakabayashi and Saito 2015; Dixon et al. 1995; Peng et al. 2017. Moreover, glycosyl-transferase-mediated glycosylation is crucial for modulating the stability, bioavailability, and biological activity of flavonoids, including anthocyanins (Le Roy et al. 2016). In addition, flavonoids modulate lipid fluidity by sterically hindering the diffusion of free radicals, and preventing membrane peroxidation, thereby contributing to membrane stability (Arora et al. 2000). Abiotic stresses trigger extensive changes in plant metabolite profiles, and advanced metabolomic approaches, especially untargeted metabolomics, can facilitate the identification of metabolite-based markers that bridge the gap between genotype and phenotype (Schrimpe-Rutledge et al. 2016). Previous studies have shown that signaling molecules such as serotonin and gentisic acid in leaves may contribute to NaCl tolerance in rice (Gupta and De 2017). Moreover, Zhao et al. (2014) reported that salt-tolerant genotypes generally accumulate lower levels of organic acids under salt stress conditions. The wild rice *O. coarctata* exhibited increased accumulation of CP47 protein, CRT/DRE-binding protein, and alcohol dehydrogenase1 under 400 mM salinity (Tamanna et al. 2024).

Rice also produces commercially valuable aromatic compounds and bioactive metabolites, including antioxidants. Identifying unknown compounds and characterizing their biological activities represent important initial steps toward uncovering the vast reservoir of valuable metabolites within the rice metabolome. Structural elucidation of these secondary metabolites is commonly achieved using spectroscopic techniques such as UV–visible spectroscopy, Fourier transform infrared spectroscopy (FTIR), mass spectrometry (MS), and nuclear magnetic resonance (NMR). In addition, advances in compound separation methods, particularly chromatography, together with mass spectrometry-based analysis of molecular ions by mass-to-charge ratio, have enabled the generation of high-quality digital datasets for comprehensive metabolomic studies. More than 1,000 metabolites, including primary and secondary metabolites, have been identified through three major plant metabolite databases, namely RiceCyc (Dharmawardhana et al. 2013), OryzaCyc, and AraCyc within the Plant Metabolic Network. Among these identified metabolites, oryzalexins and momilactones are recognized as rice-specific compounds (Kusano et al. 2015; Akatsuka et al. 1985; Kato et al. 1993; Kato et al. 1994; Kato-Noguchi and Peters 2013). Additionally, volatile phytohormones such as salicylate, jasmonate, and ethylene are also produced in rice in response to abiotic stress conditions (Iwai et al. 2006; Hattori et al. 2009; Tong et al. 2012).

Lipid remodeling plays a central role in plant adaptations to abiotic and biotic stresses (Hartel et al. 2000; Benning and Ohta 2005). Previous studies have shown that lipid remodeling promotes the replacement of membrane phospholipids with non-phosphorus glycerolipids under abiotic stress conditions, including salinity stress. Glucuronosyl diacylglycerol, a novel plant lipid that accumulates under phosphorus-limited conditions in rice and *Arabidopsis*, contributes to the maintenance of plant physiology during phosphorus deprivation (Okazaki et al. 2013). In addition, fatty acids such as linoleate and linolenate serve as precursors for jasmonic acid biosynthesis and for the formation of oxidized lipids. Notably, both processes are critical for plant responses to abiotic stress (Weber et al. 2002; Yan et al. 2022).

To elucidate the metabolic basis of salinity tolerance, untargeted liquid chromatography-mass spectrometry (LC-MS) metabolomic profiling was performed using leaf tissues collected from the introgression line JN100 (JN), its salt-susceptible recurrent parent Jupiter (JU), and the salt-tolerant donor parent Nona Bokra (NB) under control and salt stress conditions (12 dS/m EC). This study investigates how distinct metabolic signatures in the introgression line JN100 contribute to salinity tolerance and shape the underlying mechanisms and adaptive responses to salt stress.

## Material and Methods

### Plant material

Previously, our laboratory developed an introgression line (IL) population (BC_3_F_4_ generation) from a cross between Jupiter (JU) and Nona Bokra (NB) with JU as the recurrent parent (RP). ILs carrying NB genomic segments in the JU background were developed to map QTLs associated with various salinity-related traits at the seedling stage (Puram et al. 2017). NB, a salt-tolerant but low-yielding landrace with several domestication-related traits, such as tall plant stature, seed dormancy, seed shattering, and photosensitivity, was used as the donor parent. Jupiter, a medium-grain, high-yielding cultivar released by the Louisiana State University Agricultural Center (LSU AgCenter), USA, is susceptible to salinity stress and served as the recurrent parent (Sha et al. 2006). Based on seedling-stage performance under saline stress, the IL JN100 was selected because of its high level of salinity tolerance, which was comparable to that of NB (Chaudhary et al. 2025). This line was then advanced by selfing for two additional generations to fix the remaining heterozygous region.

### Salinity stress screening

Salinity treatment was imposed following the protocol described previously (Chaudhary et al. 2025). Briefly, JN, JU, and NB seedlings were grown in a greenhouse at the Central Research Station of the Louisiana State University Agricultural Center, under temperatures ranging from 28 to 30°C and under ambient light conditions. After germination in reverse osmosis (RO) water at 28°C, the seedlings were transferred to a nutrient solution and maintained until the two-leaf stage. Salt stress was then applied stepwise: seedlings were exposed to an electrical conductivity (EC) of 6 dS m ¹ for 2 days, followed by 12 dS m ¹ for 3 days, while untreated seedlings served as the control. Leaf tissues from control and salt-treated plants were harvested for metabolomic analysis.

### Sample preparation and metabolite extraction from leaf tissue

Leaf tissues from the salt-tolerant IL JN100, recurrent parent JU, and donor parent NB were used for mass spectrometry (MS) analysis. Approximately 10 mg of pulverized leaf tissue was weighed and homogenized using a pre-chilled mortar and pestle in liquid nitrogen. The powdered tissue was resuspended in 200 μl of methanol and sonicated for 10 minutes and vortexed every 2 minutes. Samples were then incubated at -20 °C for 30 minutes, followed by centrifugation at 10,000 g for 10 minutes. The resulting supernatant (∼550 μl) was transferred to a new microcentrifuge tube (MCT). To extract the organic phase, 500 μl of methyl tert-butyl ether (MTBE) was added, and the samples were vortexed and centrifuged at 10,000 g for 10 minutes. If no distinct phase separation was observed, an additional half-volume of methanol was added, followed by centrifugation at 10,000g for 10 minutes. Samples were then incubated on ice for 5 minutes to enhance complete phase separation. After centrifugation, approximately 400 μl of the upper organic layer was collected and transferred to a glass vial. The samples were incubated again at -20°C for 30 minutes and centrifuged at 10,000g for 10 minutes, after which the supernatant was transferred to a new MCT. An additional 400 μl of MTBE was then added to the pellet, and the samples were vortexed and centrifuged. The resulting MTBE layers (∼800 μl) were pooled into the same glass vial.

For lipidomic profiling, the combined organic (lipid) extract was dried under a gentle stream of nitrogen at -40 °C for 5 minutes. The remaining aqueous/methanol layer was incubated at -20°C for 30 minutes, followed by ultracentrifugation at 15,000g for 10 minutes to remove any residual MTBE. The final supernatant was then centrifuged again at 10,000 g for 10 minutes, and approximately 300 μL of the bottom polar layer was collected in a 1.5 mL MCT tube and stored at -20 °C until MS analysis. Mass spectrometry experiments for total untargeted metabolomic profiling was performed at the Louisiana State University (LSU) Mass Spectrometry Facility (MSF).

### Untargeted metabolomics using Liquid Chromatography-Mass Spectrometry

Untargeted metabolomic analysis was conducted at the LSU MSF using a Waters Synapt XS Q-TOF mass spectrometer interfaced with an Acquity Premier UPLC system. The method was designed to detect a broad range of polar metabolites rather than targeting a specific metabolite class. Metabolites were separated on a HILIC amide column using a mobile phase system consisting of acetonitrile and water supplemented with formic acid. This chromatographic method was selected to maximize the resolution and detection of small polar compounds, such as amino acids, sugars, and tricarboxylic acid cycle intermediates. Mass spectrometric data were acquired in MSE mode using the Waters all-ions fragmentation approach, without ion mobility.

### Pre-processing of the data and filtering

Untargeted LC–MS data were processed using Waters Progenesis QI v3.0. Raw files were imported and recalibrated using the accurate mass obtained from the lockspray function. Runs were aligned to correct non-linear retention time shifts, and peak picking was performed to detect features across samples. Feature annotation was performed using both MS/MS spectral matching and structure-based searches. For polar metabolites, METLIN 2019 was used as the primary MS/MS reference database. Putative identifications were scored based on fragmentation matching, mass accuracy, isotopic pattern agreement, retention-time agreement, and collision cross-section agreement. When multiple matches were obtained, the highest-scoring candidate was retained. Unmatched features were further searched against structural databases such as HMDB using theoretical fragmentation, and the same score threshold was applied. In-house-curated MS/MS databases generated on the same instrument and under the same chromatographic conditions were also used, whenever available, to improve annotation confidence.

### Metabolite identification and data processing

Metabolomics data were processed and analyzed using MetaboAnalyst (version 6.0), a web-based platform for comprehensive metabolomics data analysis, including data preprocessing, normalization, multivariate statistics, and pathway enrichment analysis. The resulting peak-intensity table was uploaded to MetaboAnalyst for log transformation and auto-scaling, followed by univariate (fold change, t-test, FDR correction) and multivariate analyses (principal component analysis, power analysis, orthogonal partial least squares discriminant analysis). Metabolite set enrichment analysis (MSEA) and metabolic pathway analysis were performed in MetaboAnalyst using species-specific *Oryza Sativa* KEGG pathways, integrating pathway enrichment and topology measures to identify significantly perturbed pathways. Pathway significance was evaluated using hypergeometric tests and topology metrics (impact scores) with default MetaboAnalyst settings. Differentially expressed metabolites identified from the MetaboAnalyst analysis were submitted as a list to the STITCH database (http://stitch.embl.de) using the batch import option to retrieve interaction data for network construction

### Metabolite set enrichment analysis

To further classify the identified metabolites into biologically meaningful subclasses, metabolite set enrichment analysis (MSEA) was carried out in MetaboAnalyst using the metabolite list generated from the JN-JU comparison under salt stress. Enrichment was assessed using over-representation analysis (ORA) with the hypergeometric test to determine whether specific metabolite subclasses were represented more frequently than expected by chance relative to the reference background. The resulting *p-values* were adjusted for multiple testing, and significantly enriched subclasses were used to support the biological interpretation of salinity-responsive metabolic changes.

### Statistical Analysis

For untargeted metabolomic datasets, statistical analyses were carried out using feature data generated in Waters Progenesis QI, in combination with the integrated EZInfo statistical package. These analyses included principal component analysis (PCA), power analysis, orthogonal partial least squares discriminant analysis (OPLS-DA), and other relevant univariate and multivariate methods. Univariate analysis was conducted using unpaired *t*-tests for pairwise comparisons between treatments within each cultivar. To reduce false positives arising from multiple comparisons, *p-*values were adjusted using the Benjamini–Hochberg false discovery rate (FDR) correction. Multivariate analyses comprised hierarchical cluster analysis (HCA), PCA, and OPLS-DA. PCA was performed on mean-centered, unit-variance-scaled data to examine the principal sources of variation in metabolic profiles. To further distinguish predefined groups (control and salt-treated samples), OPLS-DA, a supervised multivariate method, was used to maximize group separation and identify discriminant metabolites. OPLS-DA models were constructed with one predictive component and one or more orthogonal components to separate treatment-related variation from within-class variation unrelated to treatment. Variable importance in projection (VIP) scores, calculated from the OPLS-DA models, were used to identify metabolites driving group separation. Metabolites with VIP scores ≥ 1.0 were considered significant contributors to class discrimination. In addition, post hoc power analysis indicated that all statistically significant differences had power value greater than 0.8, supporting the robustness of the detected biological effects.

## Results

### Global metabolite profiling reveals genotype-specific responses to salinity stress

Multivariate analysis revealed clear metabolic differentiation among the introgression line JN100, the recurrent parent JU, and the donor parent NB under salt stress, with biological replicates clustering consistently by genotype and treatment condition. The observed sample clustering supports the robustness and reproducibility of the experimental design and further confirms that salt stress induced cultivar-specific metabolomic responses. A total of 201 DAMs with 89 upregulated and 112 downregulated metabolites were identified in the JN-JU cultivar pair under salt stress (Fig. 1).

**Figure 1.**
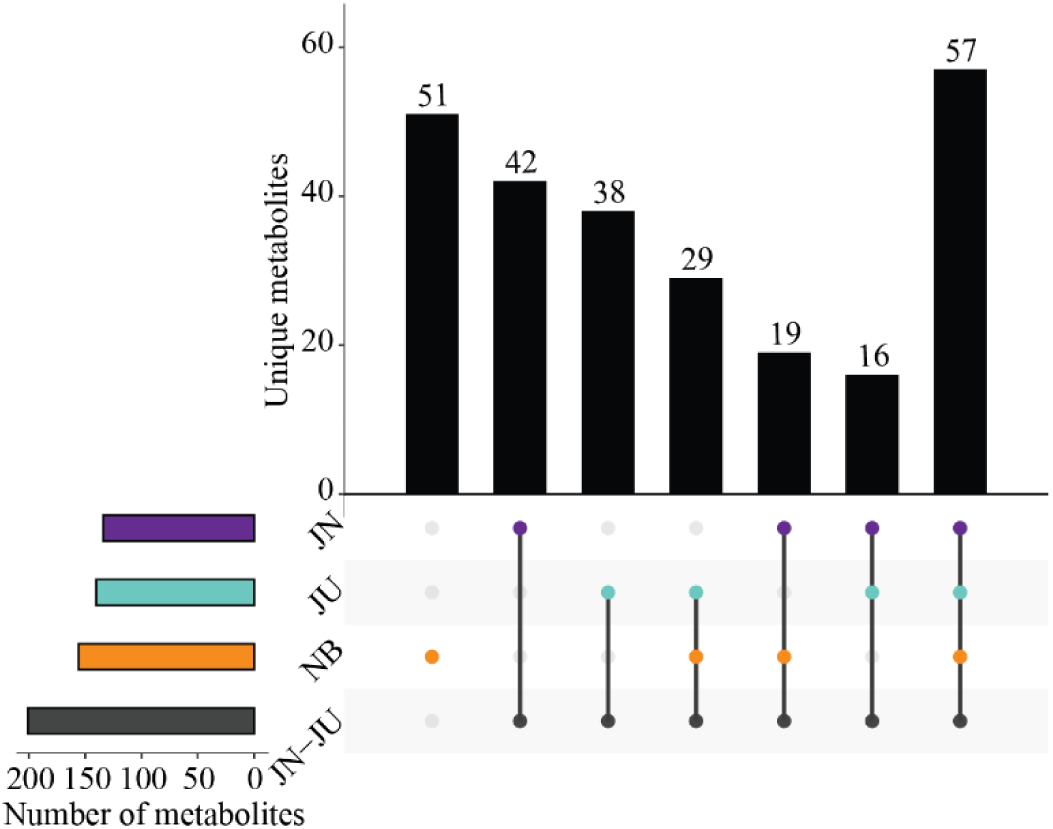
UpSet plots showing the distribution and intersections of differentially accumulated metabolites (DAMs) among JN100, Jupiter, Nona Bokra, and the JN100-Jupiter comparison under salt stress conditions. Horizontal bars indicate the total number of DAMs in each group, whereas the connected dots and vertical bars represent unique and shared metabolite sets across comparisons.

Additionally, cultivar-specific comparisons identified 134 (16 upregulated; 118 downregulated), 140 (19 upregulated; 121 downregulated), and 156 (16 upregulated; 140 downregulated) DAMs in JN, JU, and NB, respectively, under salinity conditions (Fig. 1). Volcano plot analysis further highlighted cultivar-specific metabolic responses to salt stress, with the introgression line JN100 showing a distinct pattern of metabolite regulation relative to its recurrent parent, Jupiter. The identified DAMs encompassed multiple metabolite classes, including amino acids (AA), carbohydrates, fatty acids, carbonyl compounds, amines, alcohols, dicarboxylic acids, purines, pyrimidines, benzoic acid, fatty alcohols, benzoyl derivatives, and hydroxy acids (Table 1 and Supplemental Table 1). To further categorize these salinity-responsive metabolites, metabolic set enrichment analysis (MSEA) was conducted for the JN-JU comparison. This analysis identified significantly enriched metabolite subclasses, indicating coordinated perturbation of chemically and functionally related compounds (Table 1). Among them, amino acids represented the largest group with 25 metabolites, followed by carbohydrates and fatty acids with 13 and 5 metabolites, respectively, while the remaining subclasses each contained 1 to 5 metabolites (Supplemental Table 1).

**Table 1.** Metabolite set enrichment analysis (MSEA) of metabolite subclasses identified in the JN-JU comparison under salt stress conditions.

| Pathway | Metabolite |
| --- | --- |
| Amino acids, peptides, and analogues | beta-Alanine; Glutathione; Glutamic acid; L-Tyrosine; L-Asparagine; Histidine; Serine; L-Aspartic acid; Ornithine; L-Arginine; Leucine; L-Pipecolic acid; Citrulline; S-Lactoyl glutathione; N-Acetyl-L-glutamic acid; N-Methyl-a-aminoisobutyric acid; Oxidized glutathione; D-Serine; D-Proline; D-Glutamine; D-Alanyl-D-alanine; Gabapentin; D-Pipecolic acid; L-Aspartate-semialdehyde; cis-4-Hydroxy-D-proline |
| Carbohydrates and carbohydrate conjugates | Glycerol; Glyceric acid; Sucrose; Ribitol; Gluconic acid; Maltotriose; Glucose 6-phosphate; L-Ribulose; Stachyose; 1-Kestose; Melezitose; Sedoheptulose 7-phosphate; D-Arabinono-1,4-lactone |
| Fatty acids and conjugates | 3-Hydroxymethylglutaric acid; Suberic acid; Traumatic acid; Tiglic acid; 8-Amino-7-oxononanoic acid |
| Carbonyl compounds | Diacetyl; 2-Hexenal; 2-Furancarboxaldehyde; 4-Acetyl-1-methylcyclohexene; 3-Ketosphingosine |
| Amines | Putrescine; Histidinal; 1-Butylamine; Gaboxadol |
| Alcohols and polyols | myo-Inositol; Shikimic acid; Acetone cyanohydrin |
| Dicarboxylic acids and derivatives | Maleic acid; Succinic acid; Malonic acid |
| Purines and purine derivatives | Adenine; Hypoxanthine |
| Purine ribonucleotides | Adenosine monophosphate; Guanosine diphosphate |
| Pyrimidines and pyrimidine derivatives | 3-Methylcytosine; 4-Amino-5-hydroxymethyl-2-methylpyrimidine |
| Benzoic acids and derivatives | 4-Hydroxybenzoic acid; Salicylic acid |
| Fatty alcohols | 1-Hexanol; 3-Hexen-1-ol |
| Benzoyl derivatives | 4-Methoxybenzaldehyde; 2,3,6-Trimethylbenzaldehyde |
| Alpha hydroxy acids and derivatives | Lactic acid |
| Beta hydroxy acids and derivatives | D-Malic acid |

### Multivariate analysis identifies key discriminatory metabolites under salinity

Orthogonal Partial Least Squares Discriminant Analysis (OPLS-DA) clearly separated the cultivars into two clusters corresponding to the control and salt stress treatments (Fig. 2A-B and Supplemental Figure2 A-C). The OPLS-DA score plot showed clear separation between the JN-JU pair and among the genotypes JN, JU, and NB under salt stress, highlighting genotype-specific metabolomic reprogramming in response to salinity. In the JN-JU pair, the predictive T-score explained 79.7% and 91.2% of the variance contributing to class separation under control and salt stress conditions, respectively. In contrast, the orthogonal T-score accounted for 11.8% and 3.8% of the variation unrelated to class discrimination under control and salt stress conditions, respectively (Fig. 2A-B). Moreover, the tight clustering of biological replicates within each group indicated high experimental consistency and strong model reliability.

**Figure 2.**
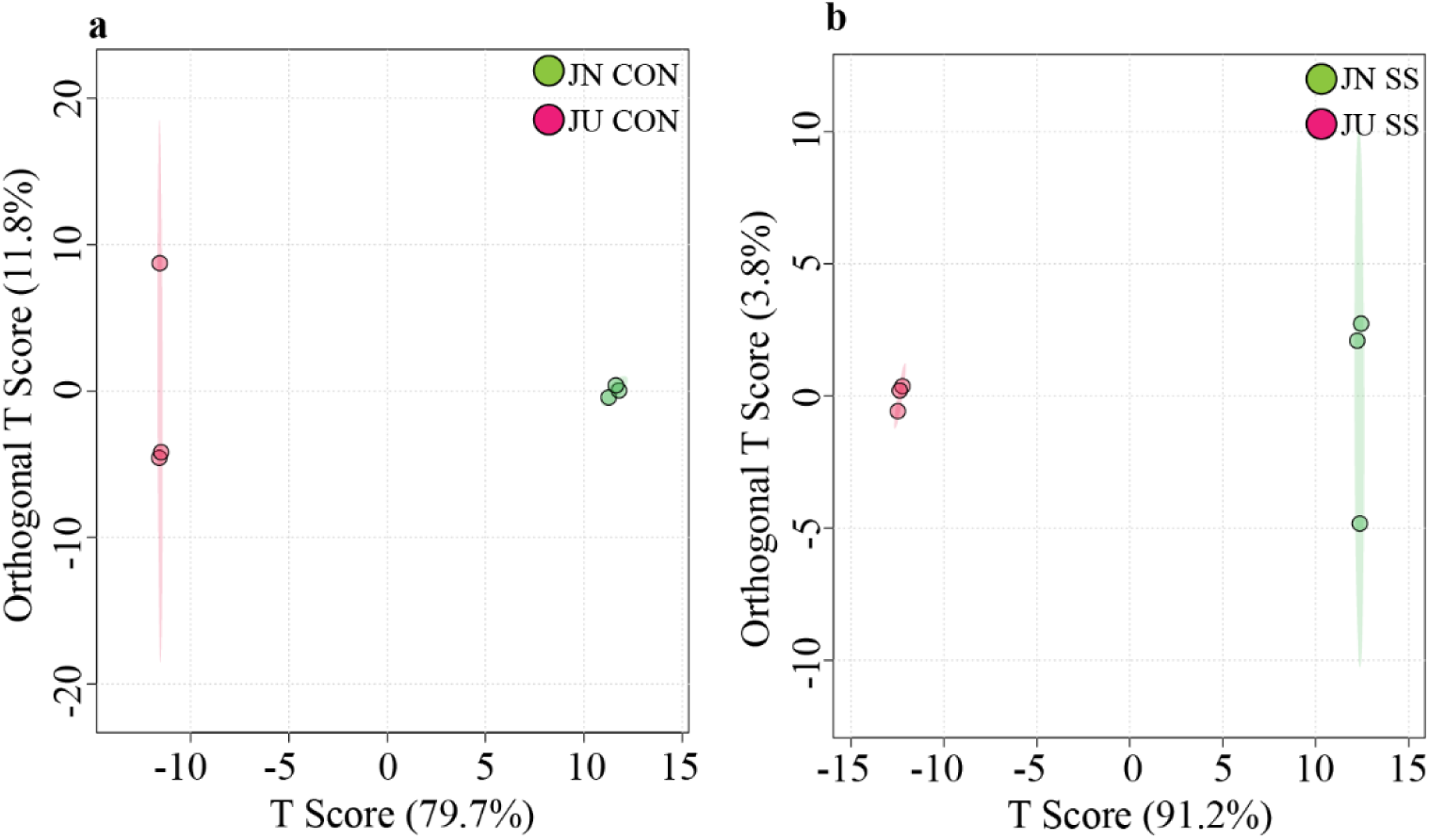
Orthogonal Partial Least Squares Discriminant Analysis (OPLS-DA) plots showing metabolic separation between JN100 and Jupiter under (A) control and (B) salt stress conditions.

Furthermore, VIP scores were used to assess the contribution of individual metabolites to group separation in the multivariate model. Metabolites with VIP scores greater than 1.0 were considered important contributors, whereas scores below 1.0 were considered less relevant in discriminating among groups. Dot plots were generated to visualize the top 15 metabolites with a VIP score greater than 1.0 across all pairwise comparisons, including JN-JU under control and salt stress conditions, as well as within the genotypes JN, JU, and NB under salt stress (Figure 3 and Supplemental Figure 1 D-F). Metabolites such as D-arabinono-1,4-lactone, violanthin, L-methionine-S-oxide, ribitol, lysine, rutin, glutamine, pantothenic acid, quinic acid, and lactulose were exclusively enriched in the JN-JU pair under salt stress (Figure 3). In contrast, berberine, 2,3,6-trimethylbenzaldehyde, glycitin, histidinal, salicylic acid, ribitol, serine, 3’-adenosine monophosphate (3′-AMP), and L-arginine were uniquely enhanced in the introgression line JN under salt stress. The salt-susceptible parent JU showed exclusive enrichment of L-asparagine, pantothenic acid, and glutamine, whereas the salt-tolerant cultivar NB showed specific enrichment of N5-methylglutamate, DL-propargylglycine, and citrulline (Supplemental Figure 1 D-F). All of these metabolites had VIP scores greater than 1, underscoring their significant contribution to genotype discrimination under salt stress (Supplemental Figure 1 D-F).

**Figure 3.**
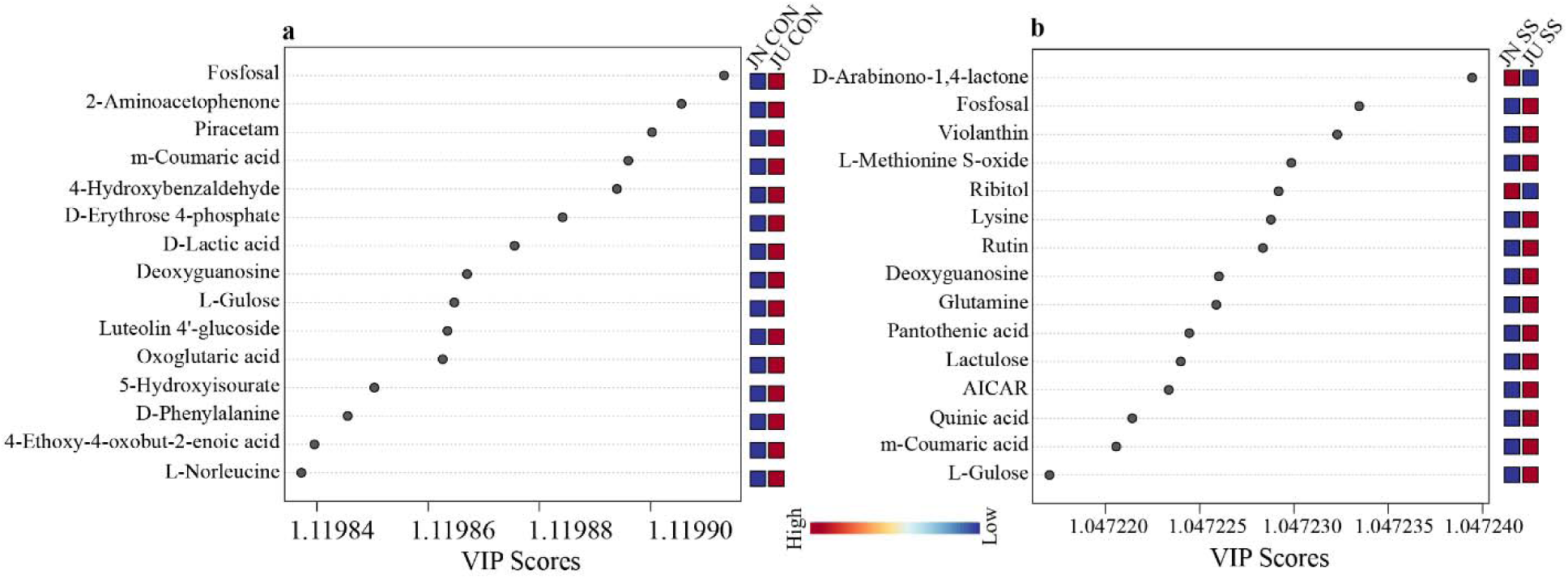
Variable importance in projection (VIP) scores showing the top discriminatory metabolites across JN100 and Jupiter under (A) control and (B) salt stress conditions. VIP scores derived from the OPLS-DA model showing the top metabolites contributing to the separation between JN100 and Jupiter under control and salt stress conditions. Metabolites with VIP scores > 1.0 were considered significant contributors to group discrimination.

### Differentially accumulated metabolites across JN100 and Jupiter under salt stress

The identified DAMs were associated with biological processes related to reactive oxygen species (ROS) signaling, oxidative stress mitigation, flavonoid and polyphenol metabolism (including flavones and flavonols), sugars and sugar alcohols, organic acids, amino acid metabolism, and phytohormonal regulation, notably jasmonate (JA) and abscisic acid (ABA)-mediated pathways (Fig. 4 and Supplemental Figure 2 A-C). These metabolite changes provide important insights into the molecular mechanisms underlying salt stress adaptation in rice. Salt stress induced distinct metabolic reprogramming in the JN-JU pair, as reflected by the differential accumulation of key metabolites associated with stress adaptation. Metabolites such as glycitin, D-arabinono-1,4-lactone (AraL), and ribitol showed increased accumulation under salt stress, whereas allantoin, violanthin, L-methionine S-oxide (MetO), lysine, and rutin, showed reduced accumulation (Fig. 4). Collectively, these DAMs appear to play major role in ROS scavenging, redox regulation, and osmotic adjustment under saline environments (Fig. 4).

**Figure 4.**
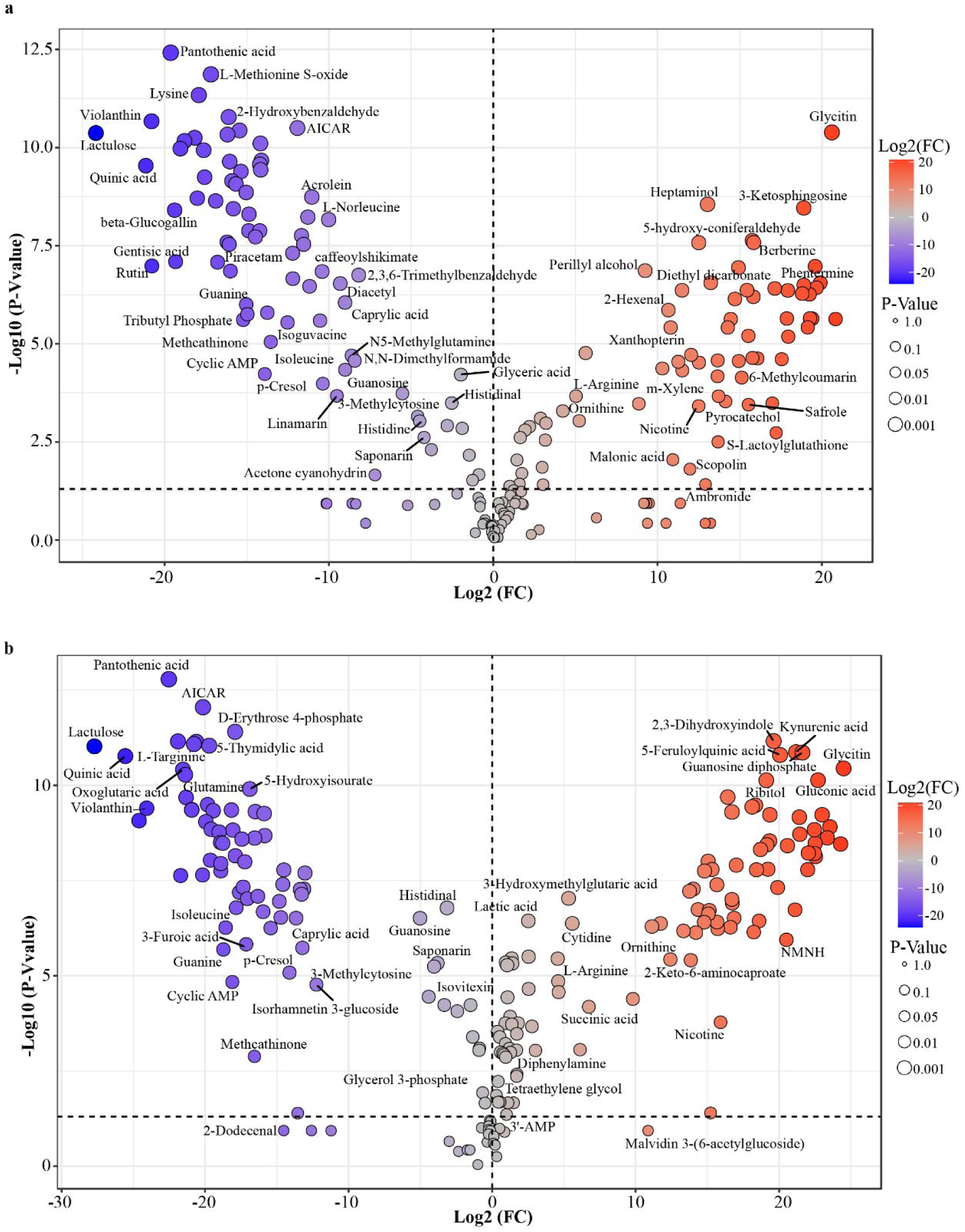
Volcano plot of differentially accumulated metabolites in the JN100-Jupiter comparison under (A) control and (B) salt stress. Each point represents a metabolite plotted by log fold change and -log adjusted *p* value. Red indicates significantly upregulated metabolites, and purple indicates significantly downregulated metabolites in JN100 relative to Jupiter under salt stress.

Allantoin, a stress-related purine metabolite, showed reduced accumulation under both control and salt stress conditions in the JN-JU pair (Fig. 4). In contrast, the isoflavone glycitin showed markedly increased accumulation, with approximately 20-fold higher level under control and nearly 25-fold higher level under salt stress in the JN-JU pair (Fig. 4 and Supplemental Table 2). Notably, glycitin was undetectable in JU when the cultivars were compared individually under salinity, indicating a genotype-specific response. In addition, AraL, a precursor of ascorbic acid, accumulated significantly in the JN-JU pair, with 23-fold and 19-fold increases under salt stress and control, respectively, and a VIP score greater than 1, placing it among the top 15 most enriched metabolites under salt stress. Similarly, ribitol showed significant accumulation, with 14-fold and 19-fold increases under control and salt stress conditions, respectively (Supplemental Table 2). Ribitol also had a VIP score greater than 1, underscoring its importance in distinguishing the metabolic profile of the salt-tolerant introgression line JN from that of its recurrent parent JU.

In contrast, several metabolites were downregulated, including violanthin, MetO, lysine, and rutin. Violanthin, a flavonoid glycoside belonging to the flavones/flavonols class, decreased by 24-fold under salt stress and 20-fold under control conditions, with VIP scores greater than 1 in both cases. MetO, a key intermediate in methionine and cysteine metabolism and a substrate for methionine sulfoxide reductases (MSRs), also showed a marked reduction, decreasing by a 17-fold under control conditions and 20-fold under salt stress, suggesting its possible conversion through MSR activity to support redox homeostasis. Likewise, lysine exhibited reduced accumulation in the JN-JU pair, with a 21-fold decrease under salt stress and a 17-fold decrease under control conditions. Notably, a significant VIP score greater than 1 was observed only under salt stress, identifying lysine as a significant DAM (Fig. 3 and Fig. 4). Similarly, rutin, a flavanol, was reduced accumulation in the JN-JU pair under both control and salt stress conditions. However, the decrease was more pronounced under salt stress, with a 24-fold decrease and a VIP score greater than 1, suggesting that rutin is also a significant DAM. (Fig. 3 and Fig. 4). Together, these metabolite patterns revealed a coordinated shift in redox-active compounds, antioxidant metabolism, amino acid turnover, and secondary metabolite biosynthesis, which may collectively contribute to salinity adaptation in rice.

### Adaptive metabolic plasticity confers salinity tolerance in rice

KEGG (Kyoto Encyclopedia of Genes and Genomes) pathway enrichment and topology analyses were performed in MetaboAnalyst using the pathway analysis function for significantly altered metabolites identified in the JN-JU pair, as well as in the individual genotypes JN, JU, and NB under both control and salt stress conditions. KEGG pathway analysis between the salt-tolerant introgression line JN and its recurrent, salt-susceptible parent JU under salt stress revealed several key differences in metabolic pathway impact, underscoring the distinct physiological responses conferred by the introgressed genomic regions (Table 1). Among the most significantly enriched pathways in the JN-JU pair, greatest impacts were observed for arginine biosynthesis and alanine, aspartate, and glutamate metabolism pathways. These pathways are central to nitrogen assimilation and osmotic adjustment, suggesting that tighter regulation of metabolites involved in these processes in JN may contribute to its greater salinity tolerance relative to JU. Further, glutathione metabolism, which is involved in antioxidant defense and ROS scavenging, glyoxylate and dicarboxylate metabolism, and pentose phosphate pathway (PPP), were enriched in the JN-JU pair under salt stress. Purine metabolism and glycine, serine and threonine metabolism also showed significant pathway impact, highlighting the reprogramming of nitrogen rich metabolites in JN (Fig. 5A). The introgression line JN showed strong enrichment of histidine metabolism, which emerged as the most significantly impacted pathway, followed by the PPP and phenylalanine; tyrosine, and tryptophan biosynthesis, indicating substantial changes in amino acid and redox-related metabolic processes (Fig. 5B). In the recurrent parent JU, arginine biosynthesis; alanine, aspartate, and glutamate metabolism exhibited the highest pathway impacts, suggesting major changes in nitrogen and amino acid metabolism. These were followed by isoquinoline alkaloid biosynthesis; butanoate metabolism; purine metabolism; glyoxylate and dicarboxylate metabolism (Fig. 5C). Moreover, the donor parent NB exhibited the highest pathway impact for arginine biosynthesis, glutathione metabolism, and arginine and proline metabolism, highlighting enhanced antioxidant and osmoprotective responses. Additional enrichment of alanine, aspartate, and glutamate metabolism, along with galactose metabolism, further supports the metabolic adjustments in NB under salt stress (Fig. 5D).

**Figure 5.**
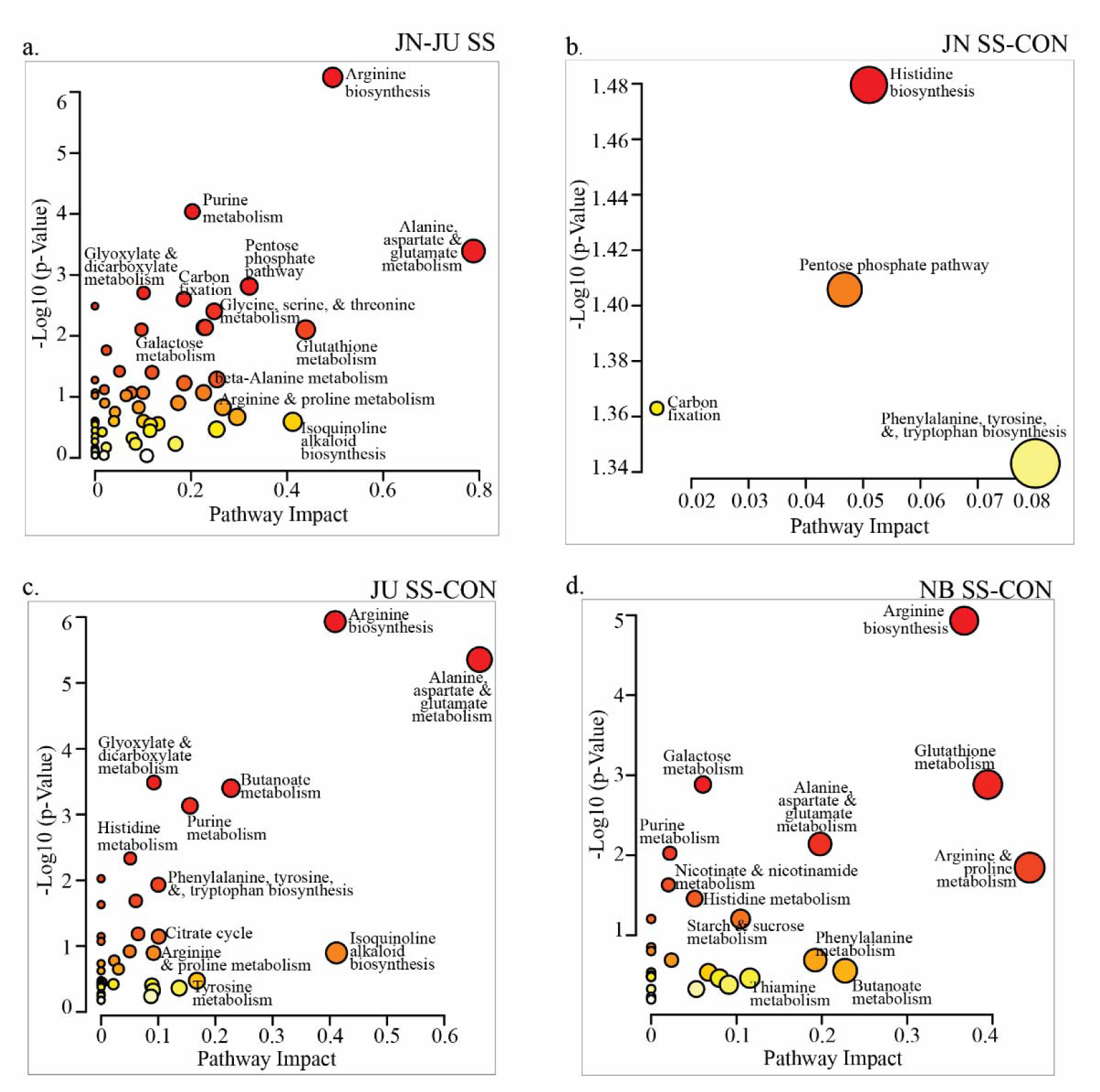
Scatter plot of enriched KEGG metabolic pathways in response to salinity stress. Pathway analysis was performed using significantly altered metabolites from the (A) JN-JU comparison under salt stress (SS), (B) JN100 SS-CON, (C) Jupiter under SS and CON (JU SS-CON), and (D) Nona Bokra under SS and CON (NB SS-CON) comparisons. In each panel, the x-axis indicates pathway impact, whereas the y-axis represents -log□□ (*p-value*) for pathway enrichment. Circle size reflects the relative pathway impact, and color intensity corresponds to enrichment significance.

Additionally, these findings are consistent with the dot plot analysis, which also showed significant enrichment of amino acid–related pathways in the JN-JU pair under salt stress, including arginine biosynthesis; alanine, aspartate, and glutamate metabolism; D-amino acid metabolism; and glycine, serine, and threonine metabolism (Fig. 6). Likewise, the enrichment of purine metabolism, the pentose phosphate pathway, carbon fixation, and glyoxylate and dicarboxylate metabolism further indicates that salinity tolerance in JN is associated with coordinated metabolic adjustments involving nitrogen metabolism, energy balance, and oxidative stress mitigation.

**Figure 6.**
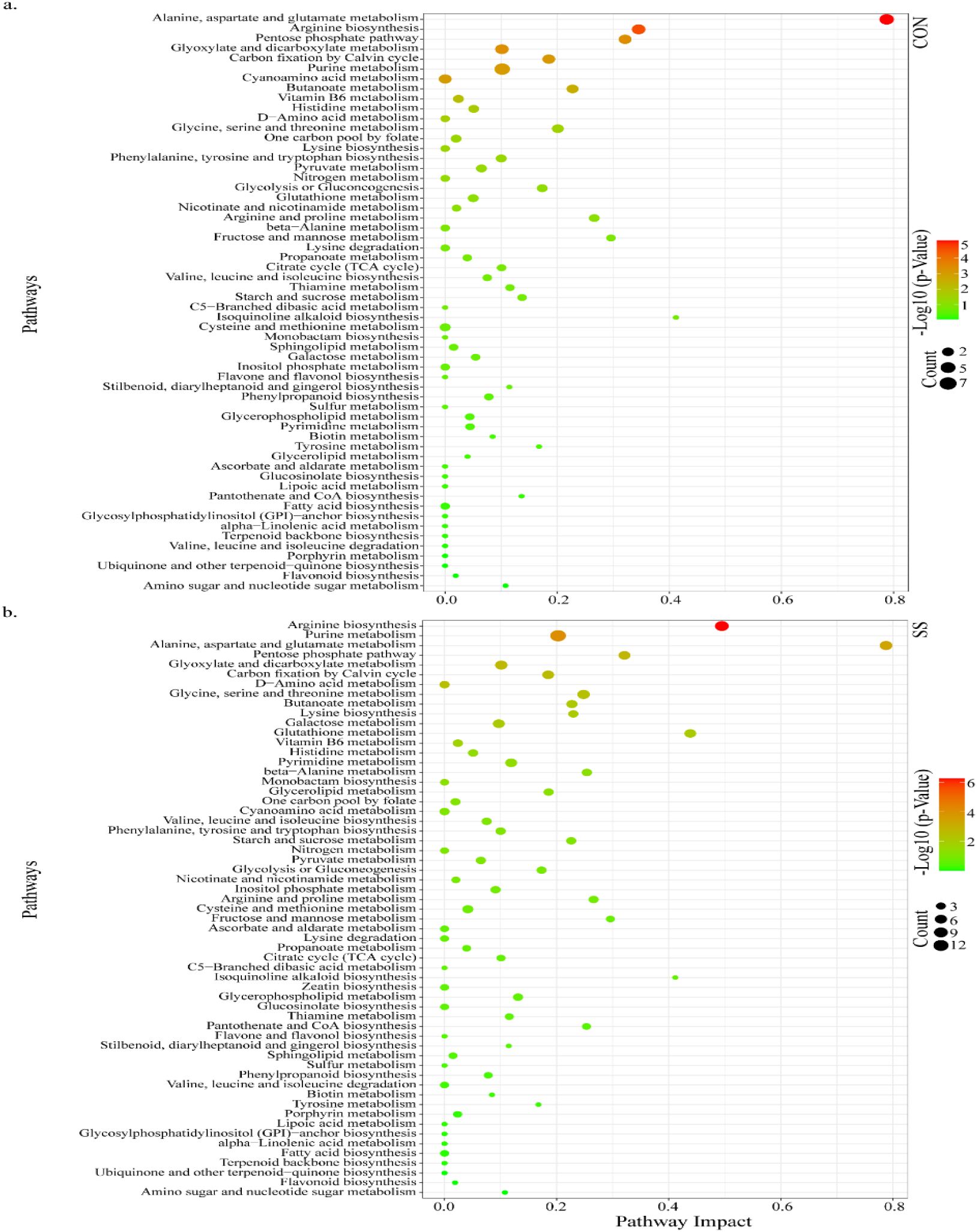
Enriched metabolic pathways in the JN-JU comparison under control and salt stress conditions. Dot plot visualization of pathway enrichment analysis based on significantly altered metabolites identified in the JN-JU pair under (A) control (CON) and (B) salt stress (SS) conditions. In each panel, the x-axis indicates pathway impact, whereas the y-axis lists the enriched pathways. Dot color represents the enrichment significance (-log□□ *p*-value), and dot size indicates the number of metabolites associated with each pathway.

Collectively, these results suggest that amino acid metabolism, redox homeostasis, and carbon–nitrogen metabolic balance are key biological responses to salinity stress.

### Joint pathway analysis

To gain an integrated view of the molecular response, the differentially expressed genes (DEGs) identified from the JN100 transcriptome dataset (Chaudhary et al. 2025) were integrated with the metabolomic profiles for joint pathway analysis.

Joint pathway analysis revealed that several pathways were significantly enriched in the JN-JU pair under salt stress, including arginine biosynthesis; glycine, serine, and threonine metabolism; glutathione metabolism; D-amino acid metabolism; and purine metabolism. These enriched pathways suggest enhanced nitrogen recycling, osmotic adjustment, and reactive oxygen species (ROS) scavenging in response to salinity stress (Fig. 7). In contrast, pathways such as aminoacyl-tRNA biosynthesis; alanine, aspartate, and glutamate metabolism; nitrogen metabolism; and carbon fixation were consistently enriched under both control and salt stress conditions, indicating that they represent core metabolic processes maintained across conditions. Notably, several pathways were exclusively enriched under salt stress, including lysine biosynthesis, linoleic acid metabolism, and fructose and mannose metabolism, indicating condition specific activation associated with stress-induced metabolic reprogramming. In addition, sulfur metabolism, pantothenate and CoA biosynthesis, and beta-alanine metabolism were selectively enriched under salt stress, further supporting the activation of genotype-specific adaptive mechanisms in JN100. By contrast, JU specifically showed enrichment of glyoxylate and dicarboxylate metabolism, suggesting a distinct metabolic adjustment in the recurrent parent under salt stress (Fig. 7).

**Figure 7.**
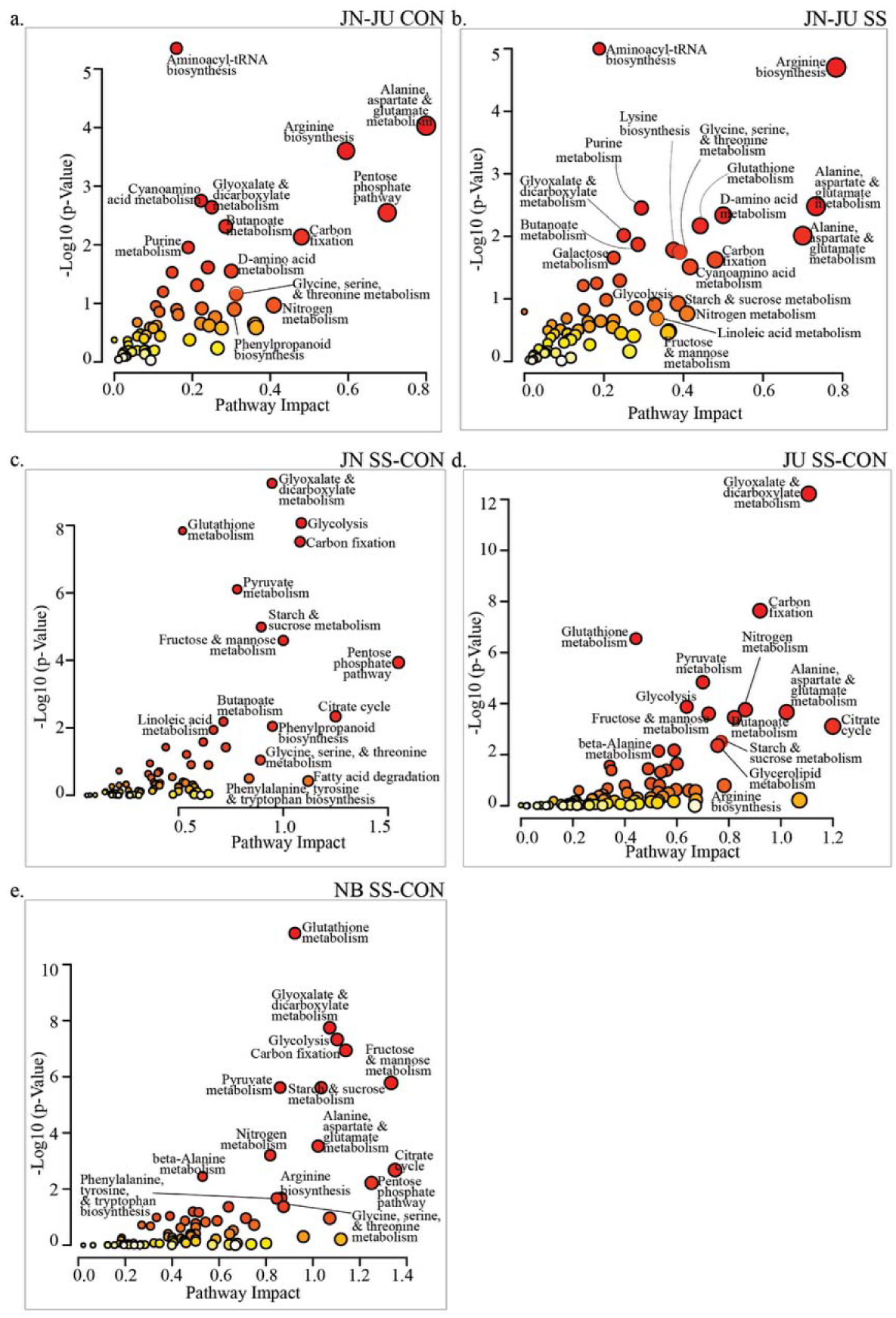
Joint pathway analysis of integrated transcriptomic and metabolomic data. Scatter plots showing enriched pathways identified by joint pathway analysis across (A) JN-JU CON, (B) JN-JU SS, (C) JN SS-CON, (D) JU SS-CON, and (E) NB SS-CON comparisons. The x-axis shows pathway impact, and the y-axis shows -log□□ (*p*-value). Circle size and color indicate pathway impact and enrichment significance, respectively.

### Integrated pathway analysis reveals coordinated transcriptomic and metabolic responses

To explore the regulatory relationship between gene expression and metabolite accumulation under salt stress, STITCH (Search Tool for Interactions of Chemicals) analysis was performed to map differentially expressed genes (DEGs) identified in the JN100 transcriptome dataset (Chaudhary et al., 2025) onto key metabolic pathways represented in the metabolomic dataset. Specifically, DEGs from the JN100 transcriptome were integrated with the corresponding metabolomic profiles of the JN-JU pair under salt stress conditions in the current study. A total of 10 DEGs overlapped with the metabolite network, of which six were functionally annotated, whereas the remaining four remain uncharacterized in rice. The annotated genes were distributed across chromosomes 1, 5, 7, 8, and 12. These included *OsDHQDT/SDH* (3-dehydroquinate dehydratase/shikimate dehydrogenase; *Os01g0375200;* Entrez ID: 4325444), *OsFd-GOGAT* (ferredoxin-dependent glutamate synthase; *Os12g0533700*; Entrez ID: 4352408), *OsALDH18B2* (aldehyde dehydrogenase 18B2, also known as proline carboxylate synthase 2; *Os01g0848200*; Entrez ID: 4344164), *OsP5CS1* (proline carboxylate synthase 1; *Os05g0455500*; Entrez ID: 4324853), phenylalanyl-tRNA synthetase (*Os12g0533700*; Entrez ID: 4338979), and a pyridoxal phosphate-dependent transferase (*Os08g0245400*; Entrez ID: 4345056), all of which were mapped to the metabolites identified in this study.

*OsDHQDT/SDH* showed involvement in the shikimate pathway, and showed predicted shikimate 3-dehydrogenase (NADP) activity, linking it to aromatic amino acid biosynthesis and secondary metabolite production (Fig. 8 and Table 2). In addition, *OsFd-GOGAT* was associated with the glutamate biosynthetic process, suggesting a role in nitrogen assimilation and amino acid metabolism. Notably, *OsALDH18B2* and *OsP5CS1*, both associated with proline biosynthesis, accumulation, osmotic stress tolerance, and ABA sensitivity, were mapped to the metabolomic profile under salt stress. These genes are also implicated in responses salt, drought, cold, and heat tolerance, as well as oxidative stress and osmotic adjustment, highlighting their functional relevance in the adaptive response of JN100 to salinity. Furthermore, phenylalanyl-tRNA synthetase was linked to the metabolomic profile as a key enzyme involved in phenylalanyl-tRNA aminoacylation, an essential step in protein synthesis and amino acid metabolism. A pyridoxal phosphate-dependent transferase associated with the biotin biosynthetic process was also linked to the identified metabolites (Fig. 8 and Table 2). Together, these integrated multi-omics findings underscore the involvement of amino acid biosynthesis, osmotic regulation, and co-factor metabolism in salinity stress responses, and particularly highlight the regulatory convergence of proline-related pathways at both transcriptomic and metabolomic levels.

**Figure 8.**
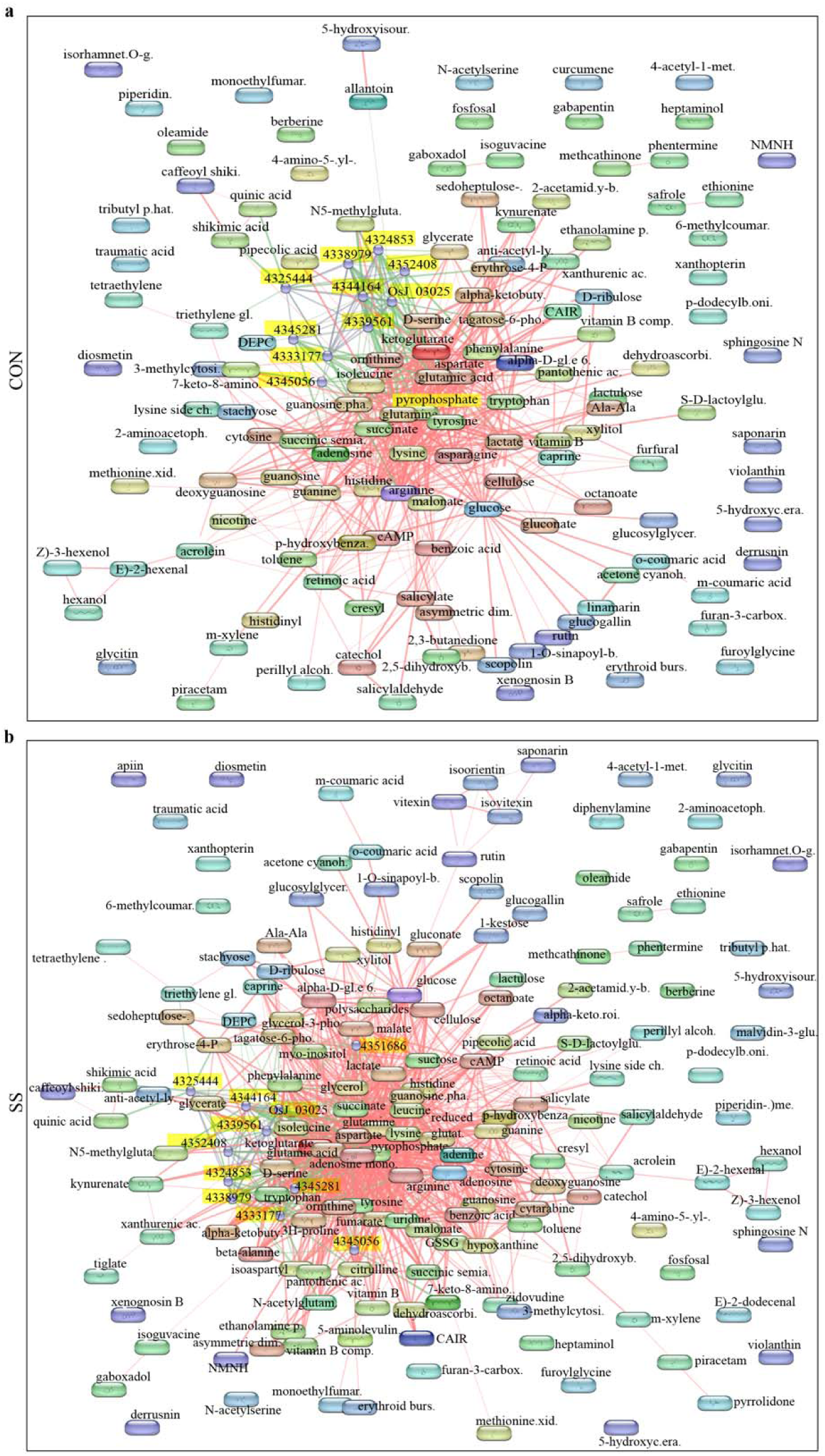
Search Tool for Interactions of Chemicals (STITCH) network analysis integrating transcriptomic and metabolomic responses in the JN-JU comparison. Gene-metabolite interaction networks constructed using STITCH for the JN-JU comparison under (A) control and (B) salt stress conditions. Differentially expressed genes (yellow nodes) were mapped to significantly altered metabolites (colored nodes) to identify functional associations between transcriptional and metabolic responses. Network edges represent predicted or database-supported interactions.

**Table 2.** Differentially expressed genes (DEGs) mapped to metabolites through Search Tool for Interactions of Chemicals (STITCH) analysis in the JN-JU comparison under salt stress. The table lists the gene ID, gene name, and functional description of DEGs.

| Gene ID | Gene Name | Description |
| --- | --- | --- |
| Os01g0375200 | *OsDHQDT/SDH* | Similar to shikimate biosynthesis protein |
| Os07g0658400 | *OsFd-GOGAT* | Ferredoxin-dependent glutamate synthase |
| Os12g0533700 | - | Phenylalanyl-tRNA synthetase |
| Os01g0848200 | *OsP5CS2* | Delta-1-pyrroline-5-carboxylate synthase 2, Salt and cold tolerance, Proline biosynthesis |
| Os05g0455500 | *OsP5CS1* | Delta-1-pyrroline-5-carboxylate synthetase 1 |
| Os08g0245400 | - | Pyridoxal phosphate-dependent transferase |

## Discussion

Understanding the physiological and metabolic basis of salt tolerance is essential for rice improvement programs because it enables identification of key traits and biomarkers associated with stress adaptation. Metabolomics offers a powerful approach for elucidating plant responses to abiotic stress by enabling the comprehensive profiling of diverse metabolites, including those generated by stress-induced metabolic reprogramming, signaling molecules involved in stress response pathways, and compounds associated with cellular acclimation under adverse conditions (Anwar et al. 2016). To further understand the metabolic basis of salt tolerance in rice, we performed leaf tissue-specific untargeted comparative metabolomic profiling of the introgression line JN100, the donor parent NonaBokra (NB), and its recurrent parent Jupiter (JU).

### Core metabolites associated with salinity adaptation

The identification of key salinity-responsive metabolites provides important insight into the metabolic mechanisms underlying salinity tolerance. In the present study, several metabolites exhibited distinct accumulation patterns under both control and salt stress conditions, indicating extensive metabolic reprogramming. These changes were associated with pathways involved in redox homeostasis, osmotic adjustment, antioxidant activity, amino acid metabolism, and stress signaling, highlighting a central role of coordinated biochemical regulation in salinity adaptation. The differential accumulation of metabolites such as glycitin, D-arabinono-1,4-lactone (AraL), violanthin, L-methionine S-oxide, ribitol, rutin, nicotinamide ribotide, lysine, and allantoin further underscores the complexity of the metabolic network underlying adaptive responses to salinity stress. Collectively, these findings suggest that salinity tolerance is supported by an integrated metabolic framework involving both constitutive and stress-induced protective mechanisms.

The strong accumulation of glycitin in JN100 relative to JU under both control and salt stress conditions indicates that this metabolite may be associated with a genotype-specific adaptive response. The fact that glycitin was undetectable Its absence in JU under salt stress when cultivars were analyzed individually further supports its selective enrichment in the introgression line. Previous studies have identified glycitin in rice metabolomic analyses associated with Pep-PEPR-mediated defense signaling (Shen et al. 2022); however, its role in salinity tolerance remains unclear. Therefore, the consistent accumulation of glycitin in JN relative to both JU and NB suggest that this isoflavone contributes to stress adaptation, potentially through antioxidant, signaling, or protective functions. These findings further indicate that glycitin may represent a previously underappreciated component of the metabolic framework associated with salinity tolerance in JN. In addition, the significant accumulation of D-arabinono-1,4-lactone (AraL) in the JN-JU comparison, particularly under salt stress, suggests a potential role in enhanced stress tolerance in JN. AraL exhibited approximately a 23-fold increase under salt stress and 19-fold increase under control, together with a VIP score greater than 1 and a ranking among the most enriched metabolites, highlighting AraL of its prominence in the JN100 metabolic response. AraL is a recognized precursor in ascorbic acid biosynthesis, and ascorbate is a central antioxidant involved in reactive oxygen species detoxification and cellular redox regulation in plants (Chen et al. 2014). Therefore, the marked enrichment of AraL in JN100 may reflect an increased capacity to support ascorbate biosynthesis, thereby improving ROS scavenging under salinity stress. Although direct evidence linking AraL to salinity adaptation is currently limited, its accumulation pattern in the present study suggests that it may contribute to maintaining redox homeostasis and protecting against oxidative damage. This response is consistent with the broader metabolic shifts observed in JN100, particularly the enrichment of antioxidant and stress-associated pathways, and supports the idea that improved oxidative stress management is a key component of salinity tolerance in this introgression line.

Violanthin, a flavonoid glycoside, exhibited reduced accumulation in the JN-JU comparison under both control and salt stress conditions, indicating differential regulation in JN100 that may be associated with stress-induced metabolic reprogramming. Violanthin is a specialized metabolite widely linked to antioxidant activity, ROS scavenging, and broader stress adaptation in plants (Radwan et al. 2026; Wu et al. 2025). Its approximately 24-fold decrease under salinity indicates that violanthin is a major discriminatory metabolite in the JN-JU comparison. Reduced violanthin accumulation may reflect depletion of flavonoid-associated antioxidant reserves under stress, or alternatively, a metabolic shift away from flavonoid glycoside accumulation toward other protective pathways that are more strongly prioritized in JN100 under salinity. This interpretation is plausible because salinity is known to trigger broad metabolic reprogramming involving antioxidant systems, redox buffering, and redistribution of metabolic resources. Moreover, ABA is a central regulator of salt-stress adaptation and is closely linked with ROS homeostasis, osmotic adjustment, and ion balance under saline conditions (Radwan et al. 2025; Wu et al. 2025). Thus, the lower violanthin levels observed here may indicate that, under salinity, JN100 preferentially supports ABA-associated protective responses and primary stress-acclimation processes rather than maintaining higher pools of this flavonoid glycoside. Although the direct role of violanthin in salinity tolerance remains unclear, its reduced accumulation in JN100 may reflect stress-induced metabolic remodeling associated with adaptive redox regulation.

Rutin, a flavonoid disaccharide derivative, exhibited a reduction in the JN-JU comparison, particularly under salt stress, suggesting that flavonoid metabolism is selectively remodeled in the introgression line JN100. Rutin has antioxidant properties, and flavonoids more broadly are known to participate in ROS scavenging and stress acclimation in plants (Wu et al. 2025). In the present study, the more pronounced decrease in rutin under salinity, together with a VIP score greater than 1 only under stress, indicates a stress-specific metabolic shift rather than constitutive suppression. This pattern may reflect the reallocation of metabolic resources away from rutin accumulation toward other protective processes that are more strongly prioritized in JN100 during salinity exposure, such as osmotic regulation, redox balancing, or central stress-signaling pathways. This interpretation is consistent with the broader metabolic signature observed in JN100, which showed enrichment of multiple pathways linked to antioxidant defense and salinity adaptation. Because exogenous rutin has been reported to mitigate salinity-associated oxidative damage in plants (Dos Santos et al. 2025; Yang et al. 2025), its depletion here may indicate active utilization or redistribution of flavonoid-associated antioxidant capacity during stress acclimation rather than a simple loss of protection.

In addition to changes in flavonoid-associated metabolism, lower accumulation of L-methionine S-oxide (MetO) in the JN-JU comparison under both control and salt stress conditions suggests altered methionine redox metabolism in JN100. MetO is an oxidized derivative of methionine and a substrate for methionine sulfoxide reductases (MSRs), which repair oxidized methionine residues through the complementary activities of MSRA and MSRB (Rey and Tarrago 2018; Guo et al. 2009). Because MSRs are closely associated with oxidative stress protection and redox homeostasis, the reduced MetO levels observed in JN100 may indicate more efficient MSR-dependent repair and regeneration of methionine, thereby limiting oxidative damage under salinity stress. Previous studies have shown that rice MSR genes, including *OsMSRA4.1,* and the overexpression of *AtMSRA4* in *Arabidopsis,* enhance tolerance to oxidative stress, further supporting the role of MSR activity in improving stress tolerance (Gustavsson et al. 2002; Guo et al. 2009).

Ribitol, a metabolite associated with redox regulation and osmoprotection, accumulated significantly in the JN-JU comparison under both control and salt stress conditions, suggesting a potential contribution to the adaptive metabolic response of JN100. Its enrichment, together with a VIP score greater than 1, highlights its importance in distinguishing the salt-tolerant introgression line from the susceptible recurrent parent. Ribitol is associated with osmoprotection and redox regulation, both of which are critical for maintaining cellular homeostasis under salinity stress. Previous studies have shown that flavonoid glycosides, including anthocyanins, accumulate under abiotic stresses such as drought, salinity, and heat, where they contribute to the alleviation of oxidative damage (Dong et al. 2020). The elevated ribitol level in JN100 therefore suggests a potential role in stress-responsive metabolic reprogramming, possibly by supporting osmotic balance and reducing oxidative damage.

Similar to ribitol, nicotinamide ribotide may contribute to the enhanced stress response of JN100 through its role in redox regulation, as it is associated with nicotinate and nicotinamide metabolism, a central pathway governing NAD(H) and NADP(H) homeostasis. These pyridine nucleotides are essential redox cofactors that support antioxidant systems, cellular energy metabolism, and ROS regulation during abiotic stress (Osterman 2009; Lu et al. 2025; Ahmad et al. 2021). Accordingly, changes in nicotinamide ribotide abundance may reflect adjustments in NAD biosynthetic and salvage pathways that maintain redox balance under salinity. Previous studies have shown that manipulation of nicotinic acid metabolism enhances drought tolerance in plants by stimulating NAD biosynthesis and strengthening stress resilience (Bashir et al. 2025; Lu et al. 2025). When considered together with the enrichment of glutathione metabolism, the pentose phosphate pathway, and other redox-associated processes in JN100, the involvement of nicotinamide ribotide further supports the view that efficient cofactor turnover and redox buffering are important components of the metabolic framework underlying salinity tolerance in this introgression line. In addition to ribitol and nicotinamide ribotide, the reduced accumulation of allantoin, a purine catabolism metabolite involved in oxidative stress regulation, suggests differential regulation of purine catabolism in JN100 relative to its recurrent parent Jupiter. Allantoin is widely recognized as a stress-associated purine metabolite with important role in oxidative stress regulation, including the reduction of endogenous H O and O levels under salinity stress (Irani and Todd 2016). In addition, allantoin has been shown to stimulate jasmonate (JA) signaling through a MYC2-regulated and ABA-dependent pathway (Takagi et al. 2016). Although JA signaling is essential for stress adaptation, excessive or prolonged activation may enhance ROS production and promote programmed cell death. Therefore, the reduced accumulation of allantoin observed in JN100 may reflect tighter regulation of ROS and JA-mediated stress signaling, thereby limiting excessive oxidative damage under salinity stress. Previous studies have shown that stress-tolerant genotypes are often characterized by more rapid and efficient activation of depolarization-activated nonselective cation channels (NSCCs), which facilitates better ROS control, whereas delayed activation through hyperpolarization-activated NSCCs can result in sustained ROS accumulation, prolonged JA signaling, and eventual cell death (Ismail et al. 2014). Taken together, the lower allantoin levels in the JN-JU comparison may represent a cultivar-specific metabolic feature associated with improved redox homeostasis and enhanced salinity tolerance in JN100.

In parallel with changes in antioxidant activity, flavonoid metabolism, and other redox-associated processes, the reduced accumulation of lysine suggests additional reprogramming of amino acid metabolism in JN100. The pronounced reduction of lysine in the JN-JU comparison, particularly under salt stress, indicates that lysine metabolism is actively remodeled in the introgression line JN100 under salt stress. Although lysine is an essential amino acid and an important component of nitrogen metabolism, its lower accumulation in JN100 may reflect enhanced utilization through stress-associated metabolic pathways rather than simple depletion.

In plants, lysine metabolism is closely linked to stress responses, and its catabolism is integrated with broader carbon–nitrogen metabolic networks (Arruda and Barreto 2020; Yang et al. 2020). Previous studies reported that amino acid metabolism contributes to nitrogen balance and stress adaptation of rice in a saline environment (Rashmi et al. 2019). The reduced lysine level in JN100 may therefore reflect its mobilization toward pathways involved in nitrogen recycling, metabolic homeostasis, or the synthesis of downstream protective compounds (Gupta and De 2017; Nazir et al. 2023). When considered together with the enrichment of arginine biosynthesis, alanine, aspartate, and glutamate metabolism, and other nitrogen-associated pathways in JN100, these findings suggest that efficient redistribution of nitrogen resources is an important component of the salinity tolerance mechanism in rice (Yang et al. 2020).

### Integrated molecular signatures of salinity response

To gain deeper insight into the molecular basis of salinity tolerance, we integrated transcriptomic and metabolomic datasets using STITCH analysis to identify gene–metabolite associations under salt stress. The overlap between DEGs and metabolite networks indicates that the metabolic changes observed in the JN-JU comparison are closely associated with transcriptional regulation of stress-responsive pathways. Notably, several annotated genes, including *OsDHQDT/SDH*, *OsFd-GOGAT*, *OsALDH18B2*, *OsP5CS1*, *phenylalanyl-tRNA synthetase*, and a *pyridoxal phosphate-dependent transferase*, were mapped to the identified metabolites, suggesting that pathways related to amino acid biosynthesis, nitrogen metabolism, and redox homeostasis are tightly coordinated during salinity adaptation.

*OsDHQDT/SDH* is associated with the shikimate pathway, and contributes to the biosynthesis of the aromatic amino acids phenylalanine, tyrosine, and tryptophan, as well as a broad range of downstream secondary metabolites (Tzin and Galili 2010). The shikimate pathway links the primary carbon metabolism with the biosynthesis of aromatic amino acids and phenylpropanoid-derived compounds. Secondary metabolites, including flavonoids and phenolics, play an important role in antioxidant defense and stress acclimation, and the association of *OsDHQDT/SDH* with these metabolites suggests that it may regulate their accumulation under salinity conditions. Moreover, phenylpropanoid metabolism is activated during abiotic stresses, including salinity, where it contributes to ROS detoxification, cellular protection, and stress adaptation (Peek and Christendat 2015; Sharma et al. 2019; Wang et al. 2017). Therefore, in the present study, *OsDHQDT/SDH* may contribute to salinity tolerance by modulating shikimate pathway flux and promoting the biosynthesis of aromatic amino acid-derived metabolites.

In addition, *OsFd-GOGAT* was mapped in the JN-JU comparison under salt stress, where it functions in the conversion of glutamine and 2-oxoglutarate into glutamate, thereby contributing to nitrogen assimilation and amino acid biosynthesis in plants. In rice, *OsFd-GOGAT* is associated with nitrogen remobilization and leaf senescence, underscoring its broader role in maintaining nitrogen metabolism and physiological homeostasis (Lee and Masclaux-Daubresse 2021; Li et al. 2022). Moreover, nitrogen metabolism is often disrupted under salinity stress, and reduced GS/GOGAT activity has been reported in rice and other crops exposed to salt stress, indicating that this pathway is highly sensitive to ionic and osmotic stress (Phan et al. 2023). The interaction of *OsFd-GOGAT* with upstream and downstream metabolites in JN100 suggests that it may contribute to salinity adaptation by sustaining glutamate production, nitrogen recycling, and amino acid homeostasis. Furthermore, alanine, aspartate, and glutamate metabolism, as well as arginine biosynthesis, were significantly enriched under salinity conditions in this study. However, further functional analyses are needed to validate the role of *OsFd-GOGAT* in rice under salt stress. Nevertheless, these findings identify it as a promising candidate gene for future studies.

Interestingly, *OsP5CS1*, involved in proline biosynthesis, was mapped onto the metabolomic network in JN100 under salinity stress. In plants, proline functions not only as an osmoprotectant but also as a stabilizer of proteins and membranes, a ROS scavenger, and a key contributor to redox buffering under abiotic stress conditions (Funck et al. 2020; Omari Alzahrani 2021). Previous studies have shown that *OsP5CS1* contributes to proline accumulation in response to stress and that ABA-mediated induction of *OsP5CS1* enhances salt tolerance by improving osmotic adjustment (Wang et al. 2017; Chen et al. 2022). *OsALDH18B2* belongs to the aldehyde dehydrogenase superfamily, which is involved in cellular homeostasis and in scavenging hydroxyl radicals through the thiol groups of Cys and Met residues. In addition, it exhibits antioxidant activity by generating NAD(P)H, which is essential for GSH regeneration (Kotchoni et al. 2010). We also observed enrichment of genes such as phenylalanyl-tRNA synthetase and a pyridoxal phosphate-dependent transferase, which are involved in translation and may therefore influence protein synthesis and amino acid utilization under salinity conditions in the JN-JU comparison (Hanson et al. 2016; Yu et al. 2025).

Collectively, these findings provide an integrated view of the metabolic and transcriptional reprogramming underlying salinity tolerance in rice. The identified metabolites, enriched pathways, and candidate genes offer mechanistic insights into the regulation of redox homeostasis, osmotic adjustment, and nitrogen metabolism under salt stress and constitute valuable molecular targets for future functional validation and rice improvement.

## Acknowledgements

We thank Dr. Fabrizio Donnarumma, LSU Mass Spectrometry Facility, for the LC-MS experiments.

## Author Contributions

PKS, MRG: supervised the experiments; PKS, MRG, CKC: designed the experiment; CKC, KA: performed the experiments, CKC, PKG: analyzed the data; CKC: wrote the paper

## Data Availability

All data reported in this paper will be shared upon request.

## Data Availability

The authors declare no competing interests.

**Supplemental Figure 1.** Orthogonal Partial Least Squares Discriminant Analysis (OPLS-DA) plots showing metabolic separation across (A) JN100, (B) Jupiter, and (C) Nona Bokra under control and salt stress conditions. Variable importance in projection (VIP) scores showing the top discriminatory metabolites across (D) JN100, (E) Jupiter, and (F) Nona Bokra under control and salt stress conditions. VIP scores derived from the OPLS-DA model showing the top metabolites contributing to the separation between JN100 and Jupiter under control and salt stress conditions. Metabolites with VIP scores > 1.0 were considered significant contributors to group discrimination.

**Supplemental Figure 2.** Volcano plot of differentially accumulated metabolites in the (A) JN100, (B) Jupiter, and (C) Nona Bokra under control and salt stress, respectively. Each point represents a metabolite plotted by log fold change and -log adjusted *p* value. Red indicates significantly upregulated metabolites, and purple indicates significantly downregulated metabolites in JN100 relative to Jupiter under salt stress.

**Supplemental Table 1.** Metabolite set enrichment analysis (MSEA) of significantly enriched pathways in the JN-JU comparison under (A) control and (B) salt stress conditions. The table summarizes the enriched metabolic pathways identified by MSEA using metabolites detected in the JN-JU comparison. For each pathway, the table provides the total number of metabolites in the reference pathway (Total), the expected number of hits by chance (Expected), the observed number of matched metabolites (Hits), the raw p value, the -log10-transformed p value [LOG10(p)], the pathway enrichment score, the Holm-adjusted p value, the false discovery rate (FDR), and the pathway impact score.

**Supplemental Table 2.** Differentially accumulated metabolites (DAMs) across the JN100, Jupiter, and Nona Bokra under salt stress conditions.

